# The ultrastructure of the starfish skeleton is correlated with mechanical stress

**DOI:** 10.1101/2024.02.07.579295

**Authors:** Raman, Susanna Labisch, Jan-Henning Dirks

## Abstract

Echinoderms and vertebrates both possess mesodermal endoskeletons. In vertebrates, the response to mechanical loads and the capacity to remodel the ultrastructure of the skeletal system are fundamental attributes of their endoskeleton. To determine whether these characteristics are also inherent in Echinoderms, we conducted a comprehensive biomechanical and morphological study on the endoskeleton of *Asterias rubens*, a representative model organism for Echinoderm skeletons. Our analysis involved high-resolution X-ray CT scans of entire individual ossicles, covering the full stereom distribution along with the attached muscles. Leveraging this data, we conducted finite element analysis to explore the correlation between mechanical loads acting on an ossicle and its corresponding stereom structure. To understand the effects of localized stress concentration, we examined stereom regions subjected to high mechanical stress and compared them to areas with lower mechanical stress. Additionally, by comparing the stereom structures of ossicles in various developmental stages, we assessed the general remodeling capacity of these ossicles. Our findings suggest that the ability to adapt to mechanical loads is a common feature of mesoderm endoskeletons within the Deuterostomia taxonomic group. However, the material remodelling may be a specific trait unique to vertebrate endoskeletons.

## Introduction

Echinoderms are marine invertebrates that possess a remarkable skeletal structure. This skeleton is histologically a mesodermal endoskeleton, however, based on its macroscopic structure a functional exoskeleton. It is an impressive example of a multipurpose tissue that serves various functions, such as structural support, locomotion, protection against predators, food gathering, feeding, and ion storage [1–3]. Given its mesodermal origin, the echinoderm endoskeleton is phylogenetically closer related to the mesodermal endoskeleton of Vertebrates [4,5].

The primary role of the Echinoderm endoskeleton is to provide the foundation for biomechanical functions, including locomotion and structural stability, by offering attachment points for muscles. In the case of most land-based vertebrates, their skeletons play a crucial role in supporting their body mass, and yet, the skeleton itself can constitute a substantial portion of their total body mass. While a heavy bone endoskeleton offers excellent support and protection, it comes with a significant cost in terms of construction, maintenance, and mobility. As a result, vertebrate endoskeletons have evolved various adaptations to optimize their mass. For instance, in vertebrate bones, the internal structure, or ultrastructure, often correlates with the level of mechanical loading. Typically, regions of bones subjected to higher mechanical stress exhibit a denser structure compared to areas with lower biomechanical stress [6]. In addition to this “static” correlation of stress and structure, which can also be found in materials like wood [7]. Interestingly, vertebrate bones can also “dynamically” remodel their structure when loading conditions change, for example during ontogenetic processes. These two abilities allow for minimum use of material while ensuring the required mechanical properties of the skeleton throughout the entire life of a vertebrate. Recently, it has been shown that Insects are likely to also show the ability to remodel their cuticle exoskeleton when subject to increased mechanical load [8].

In contrast to most vertebrate skeletons built from relatively large single bone elements, the echinoderm skeleton typically consists of numerous small, calcite structures called ossicles (Figure 1, SV 1), which are interconnected by muscles and mutable collagenous tissues (MCT) to form a complex network within the body wall [9–12]. As bones in invertebrates have a porous trabecular microstructure, a similar porous microstructure has also been observed in the echinoderm’s ossicles (SV 2). This porous microstructure exists in different geometries known as stereoms [1,2]. Different types of stereom are present within a single ossicle and are organized in complex geometrical patterns. Multiple stereom in individual ossicles might serve several different functions within the skeleton and might be subjected to different types of mechanical loads correspondingly. The stereom provides sites for different soft tissue connections, and withstand mechanical stresses arising from various activities such as feeding, locomotion, protection from predators etc. [1]. Also, the ossicles share contact surfaces with adjacent ossicles in the skeleton thereby distributing the mechanical stresses throughout the skeleton. Different stereoms might suited well for these different functions.

**Figure 1:**
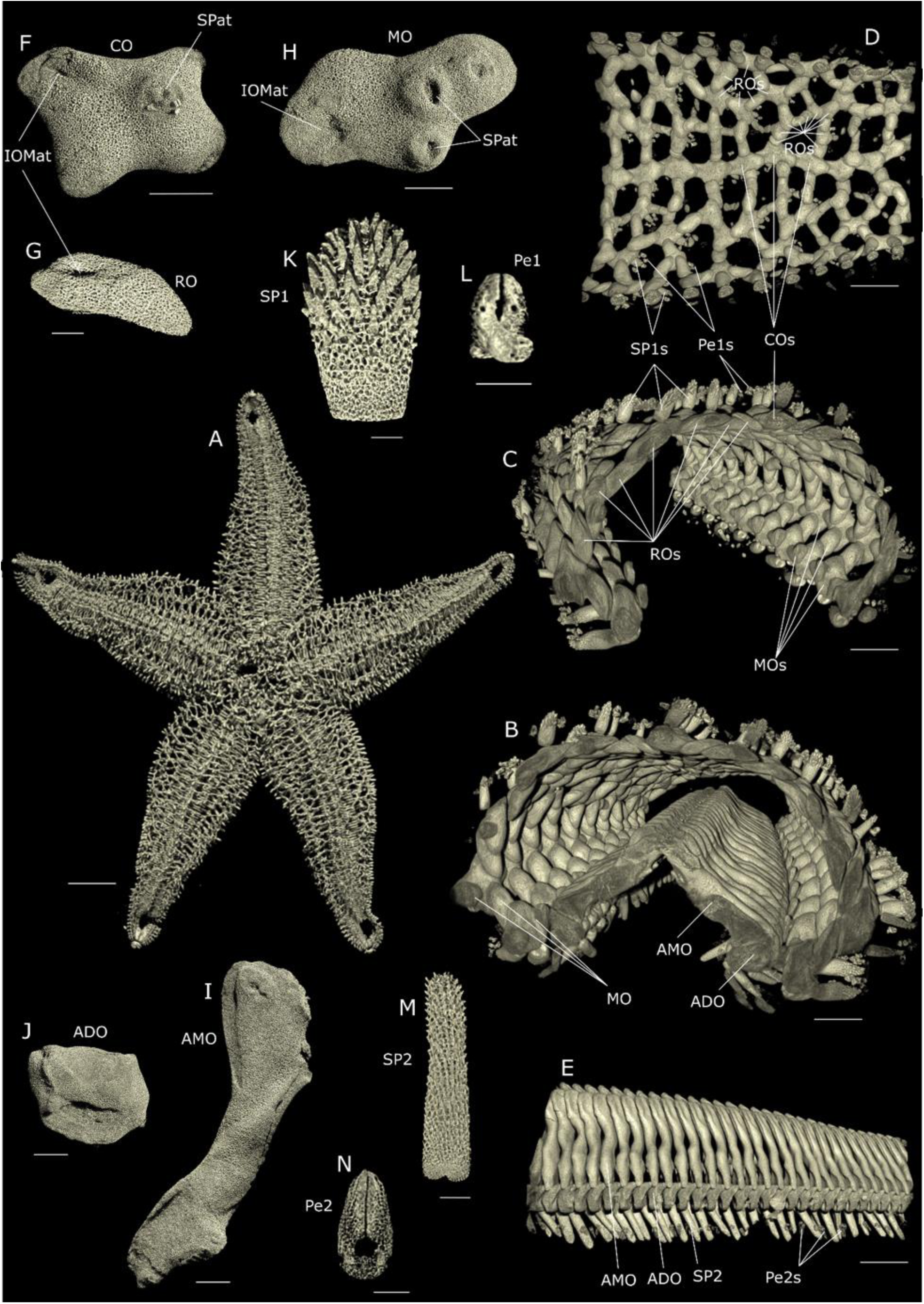
MicroCT scans of the skeleton of starfish Asterias rubens. (**A**) Overview scan of the complete starfish skeleton visualised from the aboral side (the scan was performed at 76 µm voxel size and is adapted from [35]). (**B**) Cross-sectional view of an individual ray of the starfish viewed from proximal side (the scan was performed at 13 µm voxel size and is adapted from [35]). The ossicles are arranged in two different patterns. The ambulacral (AMO) and adambulacral ossicles (ADO) form a groove shaped geometry on the oral side while the carinal (CO), marginal (MO) and reticular ossicles (RO) form a dome shape geometry on the aboral side of the ray. (**C**) Geometrical arrangement of different marginal ossicles (MO) is visualised in the absence of the ambulacral (AMO) and adambulacral ossicles (ADO) when viewed from the proximal end. This was achieved by removing the ambulacral (AMO) and adambulacral ossicles (ADO) from the 3D CT image data by defining a region of interest manually. Marginal ossicles (MO) are consistent in maintaining their shapes and geometrical arrangement throughout the length of the ray. (**D**) Geometrical arrangement of the carinal (CO) and reticular ossicles (RO) is visualised when viewed from the oral side of the ray. From left to right in the image is from the proximal to the distal end. The carinal ossicles (CO) are arranged in a row, connecting to each other in the longitudinal axis of the ray. As seen in the image, carinal ossicles (CO) not consistent with their shapes throughout the length of the ray. Carinal ossicles (CO) are connected to the reticular ossicles (RO) in the transverse axis of the ray. Reticular ossicles (RO) exist in different shapes and are arranged in ring shaped patterns. On the outside of the ray, the spines (SP1) are attached to the ossicles along with the pedicellariae (Pe1). (**E**) Geometrical arrangement of the ambulacral (AMO) and adambulacral ossicles (ADO) is visualised in the absences of all other ossicles. From left to right in the image is from the proximal to the distal end. Ambulacral (AMO) and adambulacral ossicles (ADO) have only one specific shape and the size of these ossicles decreases from proximal to distal end of the ray. Attached to the adambulacral ossicles (ADO) are the elongated spines (SP2) with pedicellariae (PE2). (**F**) An individual carinal ossicle is visualised (the scan is performed at 2 µm voxel size). Spine attachment site (SPat) and inter ossicular muscle attachment site (IOMat) is observed on the surface of the ossicle. The presence of spine attachment site (SPat) on the ossicle surface shows that this surface of ossicle must be facing towards the outside of the ray. On the ossicle surface, the spine attachment sites (SPat) are observed as depressions surrounded by an elevated boundary, while inter ossicular muscle attachment sites (IOMat) are seen as depressions on the flat surface. (**G**) An individual reticular ossicle (RO) with inter ossicular muscle attachment site (IOMat) is observed. (**H**) An individual marginal ossicle (MO) with two spine attachment sites (SPat) and one inter ossicular muscle attachment site (IOMat) is seen. Presence of spines attachment sites (SPat) shows that this surface of the ossicle must be facing outside of the ray (**I**) An individual Ambulacral ossicle (AMO) is seen from the proximal side of the ray. Ambulacral ossicles (AMO) have a very specific shape, which is constant throughout the length of the ray. (**J**) An individual Adambulacral (ADO) is seen from the proximal side of the ray. Similar to the ambulacral ossicles (AMO), they also have very specific shape, which is maintained throughout the length of the ray. (**K**) Spine (SP1), is attached to the aboral side of the ray on carinal (CO), marginal (MO) and reticular ossicles (RO). Pointed sharp edges on the top half of the spine (SP1) provide protection from predators. (**L**) Surrounding the spines (SP1), are the pedicellariae (Pe1). (**M**) Attached to the adambulacral ossicles (ADO) are the elongated shaped spines (SP2). Spines (SP2) don’t have sharp edges like spines (SP1). (**N**) Surrounding the spines (SP2) are the pedicellariae (Pe2). ((**F-N**) scans were performed at 2 µm voxel size and ossicles displaced were taken from different animals). Scale bars: (**A**) 10 mm; (**B-E**) 1 mm; (**F, H, I, J**) 500 µm; (**G, K, L, M, N**) 100 µm.

Various environmental factors play a significant role in shaping the skeletal form of an organism. These factors can include the ease of access to skeletal building materials and the mechanical stress of the surrounding environment (e.g. the requirement to carry their own body mass). These factors can also determine the evolutionary advantage of adaptations to minimize the mass of the skeleton, leading to the ability to dynamic remodel of the skeleton material during ontogenesis.

Echinoderms, in contrast to vertebrates, predominantly lead a benthic lifestyle, where the body mass poses fewer limitations on locomotion, and they expend less energy for movement [13]. Consequently, echinoderms might not have the same imperative to minimize the mass of their endoskeleton, potentially eliminating the need for dynamic remodeling. However, the availability of building materials also plays a significant role in determining the body mass of the skeleton. When there is a shortage of building materials, the necessity to recycle the existing skeleton material arises, driving the requirement for dynamic remodeling.

Earlier studies have already described the resorption and remodeling of skeletal material in sea urchins [11,14–17]. For sea urchin spines it has been shown that when the new stereom forms peripherally the thickness of the old meshwork increases by secondary deposition of minerals on previously formed meshes [1,18]. Previous studies have also shown that sea urchins possess the ability to repair their spines (of mesodermal origin) throughout their lifespan [19–21]. The repair process involves the recruitment of sclerocytes to the damaged area, followed by the deposition of new matrix and minerals to restore the integrity of the ossicle. This shows that the mesodermal endoskeletons of at least some echinoderms possess the ability to repair damage. However, so far, no systematic study has focused on analyzing the remodelling of already deposited material in the skeletal ossicles of Echinoderms.

If Echinoderms do possess the ability to remodel their ossicle structure, then the reaction to the mechanical loads and the ability to remodel is a fundamental characteristic of all mesodermal endoskeletons. If Echinoderms do not possess these two “features”, they must have developed at a later stage in osteocytic bone evolution.

In the past, the analysis of the complex ultrastructure of Echinoderm skeletons has been experimentally very challenging. These studies have often relied on 2D information obtained from scanning electron microscope (SEM) images of stereom structures. However, using limited 2D information alone, it becomes exceedingly difficult to accurately predict the distribution of porosity and mechanical stress within an ossicle. Consequently, investigating potential correlations between stereom structure and mechanical stresses was nearly impossible [22–24].

Instead, comprehensive 3D information on the entire endoskeleton structure which includes muscle attachment sites, contact sites and stereom geometries is required to investigate the correlation of morphology and biomechanics. Such data can be provided by high-resolution X-ray computed tomography (CT) which allows for non-destructive analysis of complex 3D structures. However, with most mictoCT techniques obtaining suitable 3D information on stereom patterns provides a significant experimental challenge because the range of structural thicknesses in echinoderm stereom does vary, spanning from as little as 1 µm to a maximum of 30 µm [25]. Thus, to capture the complete stereom geometry, CT scans need to be performed at a much higher resolution. With increasing resolution, however, the respective field of view decreases notably, thus less of the complex 3D structure of the stereom within the ossicle can be analyzed. As a consequence, previous studies on the 3D stereom structure could just focus on a part of the ossicle instead of the complete ossicle geometry [26–30]. Without the complete ossicle geometry, the position of muscle attachment sites and contact sites, and the 3D structure of the stereom it was not possible to accurately determine the actual mechanical loading experienced by an ossicle and thereby analyse the induced mechanical stresses.

To investigate whether Echinoderm possesses a “static” correlation of mechanical load with stereom structure and the “dynamic” ability to remodel their ossicles we thus carried out a comprehensive biomechanical and morphological study on the endoskeleton of *Asterias rubens*, a typical biological model organism for Echinoderm skeletons. We conducted high-resolution X-ray CT scans on entire individual ossicles, encompassing the full stereom distribution as well as the attached muscles. We utilized this data to carry out a finite element analysis, aiming to investigate the "static" correlation between the mechanical loads experienced by an ossicle and its corresponding stereom structure. To understand the effect of localized stress concentration, we also examined the stereom in regions experiencing localized high mechanical stress and compared it to locations with lower mechanical stress. Finally, by comparing the stereom structures of ossicles in different developmental stages, we investigated the general remodeling capability of ossicles.

## Results

### Porosity of different ossicle types

Ossicles located on different sites in a starfish arm are subjected to different types of mechanical stresses due to their location and the specific task they perform during locomotion and feeding [31–33]. For example, during feeding, the tube feet near the base of the starfish rays are used for opening shells and in turn exert direct forces on the ambulacral ossicles to which they are connected. The ossicles at the tip of the ray do not perform this function and are not subjected to such type of loading [3]. If the stereom of the ossicles is affected by the mechanical load, then different ossicle types from various locations along the ray should have variations in their stereom structure.

To test this hypothesis, we compared the average percentage porosity (APP) of all five different types of ossicles (ambulacral, adambulacral, carinal, marginal and reticular ossicles in Fig 1.), extracted from the three different locations of the starfish rays (Fig 2.). The APP describes the overall relation of material to “non material” in a given volume.

**Figure 2:**
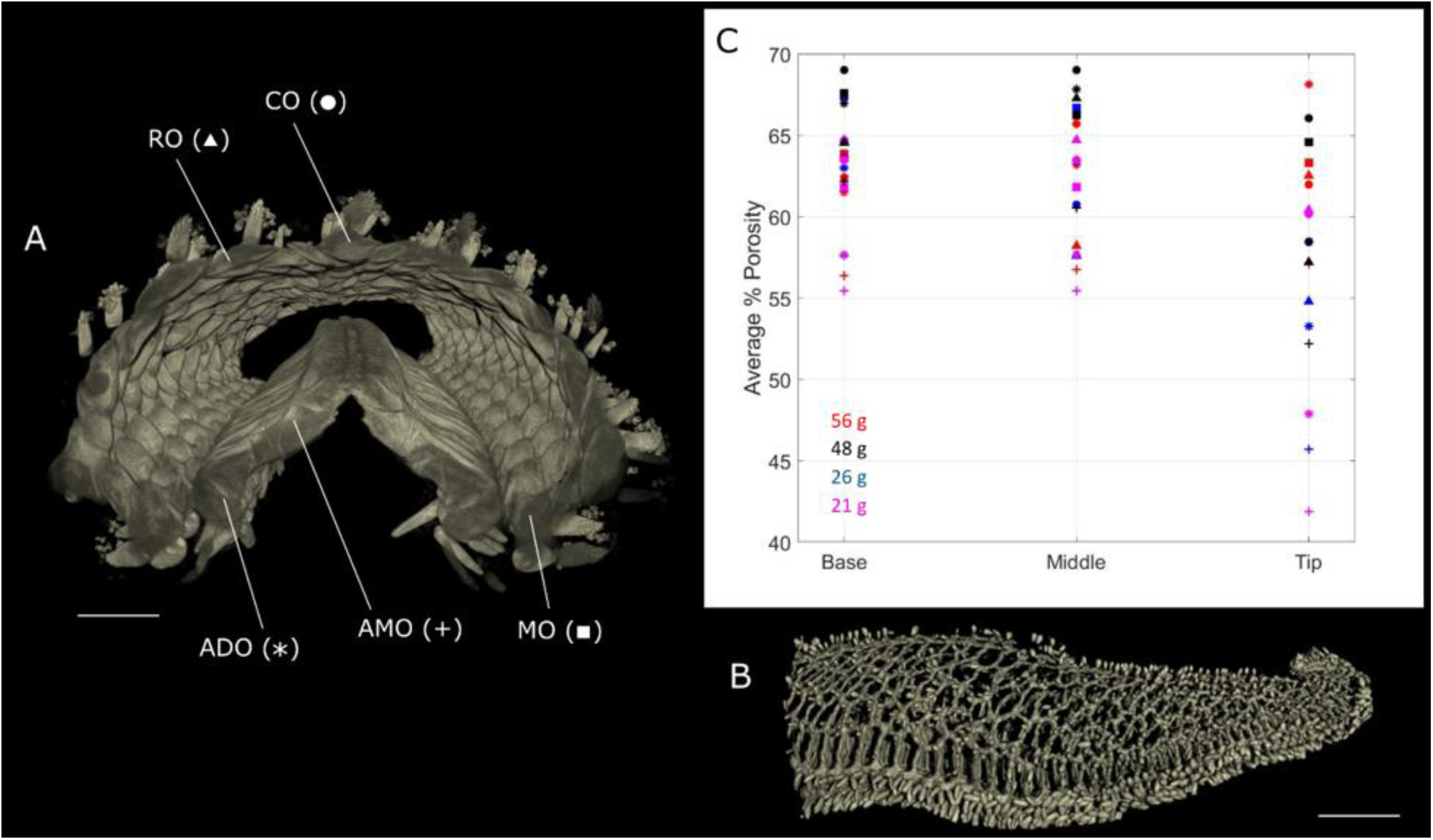
Average percentage porosity comparison of different ossicles (**A**) X-ray CT scan image of cross-sectional view of starfish ray from proximal side. Different ossicles are assigned with different symbols: Ambulacral (AMO (+)), Adambulacral (ADO (*)), Marginal (MO(□)), reticular ossicle (RO(△)) and coronal ossicle (CO(O)). (scan was performed at 13 µm voxel size and adapted from [35]) (**B**) X-ray CT scan image of an individual starfish ray. Base, middle and tip regions of the ray are aligned to the graph above, depicting that from these regions, the ossicles were extracted for average percentage porosity comparison. (the scan was performed at 50 µm voxel size) (**C**) Graph showing the comparison of average percentage porosity of different ossicle types, extracted from different locations i.e., base, middle and tip region of starfish ray. The study is performed on four different starfish with body mass ranging from (21-56) grams. Scale bars: (**A**) 1mm; (**B**) 10 mm.

Our analysis shows that both ossicle type and ontogenetic stage had a significant effect on the measured APP (Two-factor ANOVA, type: F_4_=9.799, p<0.001, stage: F_2_=7.400, p<0.001). Ossicles from the tip region of the ray showed an APP that was significantly lower than the APP of ossicles from the middle or base region (tip-middle and tip-base p<0.01, middle-base p>0.05). Across all ontogenetic stages, the APP of ambulacral ossicles was significantly lower than the APP of all other ossicles. There were no significant differences in APP between the other ossicle groups (Table 1.

**Table 1:** Summary of statistical analysis of APP difference between different types of ossicles across all ontogenetic stages. Significant differences with p>0.05 are highlighted. Across all ontogenetic stages, the APP of ambulacral ossicles (AMO) was significantly lower than the APP of all other ossicles. There were no significant differences in APP between the other ossicle groups.

|  | AMO | ADO | CO | MO |
| --- | --- | --- | --- | --- |
| ADO | <0.001 | -- | -- | -- |
| CO | <0.001 | >0.05 | -- | -- |
| MO | <0.001 | >0.05 | >0.05 | -- |
| RO | <0.01 | >0.05 | >0.05 | >0.05 |

### Stereom structure at muscle attachment sites

Inter ossicular muscles (IOMs) connect ossicles and when contracted, move the ossicles relative to each other. This movement, in turn, allows the ray to change its shape. IOMs loop around on the stereom on the surface of the ossicle and establish strong connections. When these IOMs contract, the stereom on which they are connected directly experiences the pulling force. While the stereom at the non-muscle attachment sites on the ossicle surface never experiences this type of force. Major load-carrying ossicles such as ambulacral ossicles, are connected to the biggest and highest number of inter ossicular muscles (Figure 3 (A-D, B1, B11, B12, C1)). If the stereom of the ossicle is correlated with the mechanical stresses induced by IOMs pull, then the stereom at muscle attachment sites should be different from that of non-muscle attachment sites.

**Figure 3:**
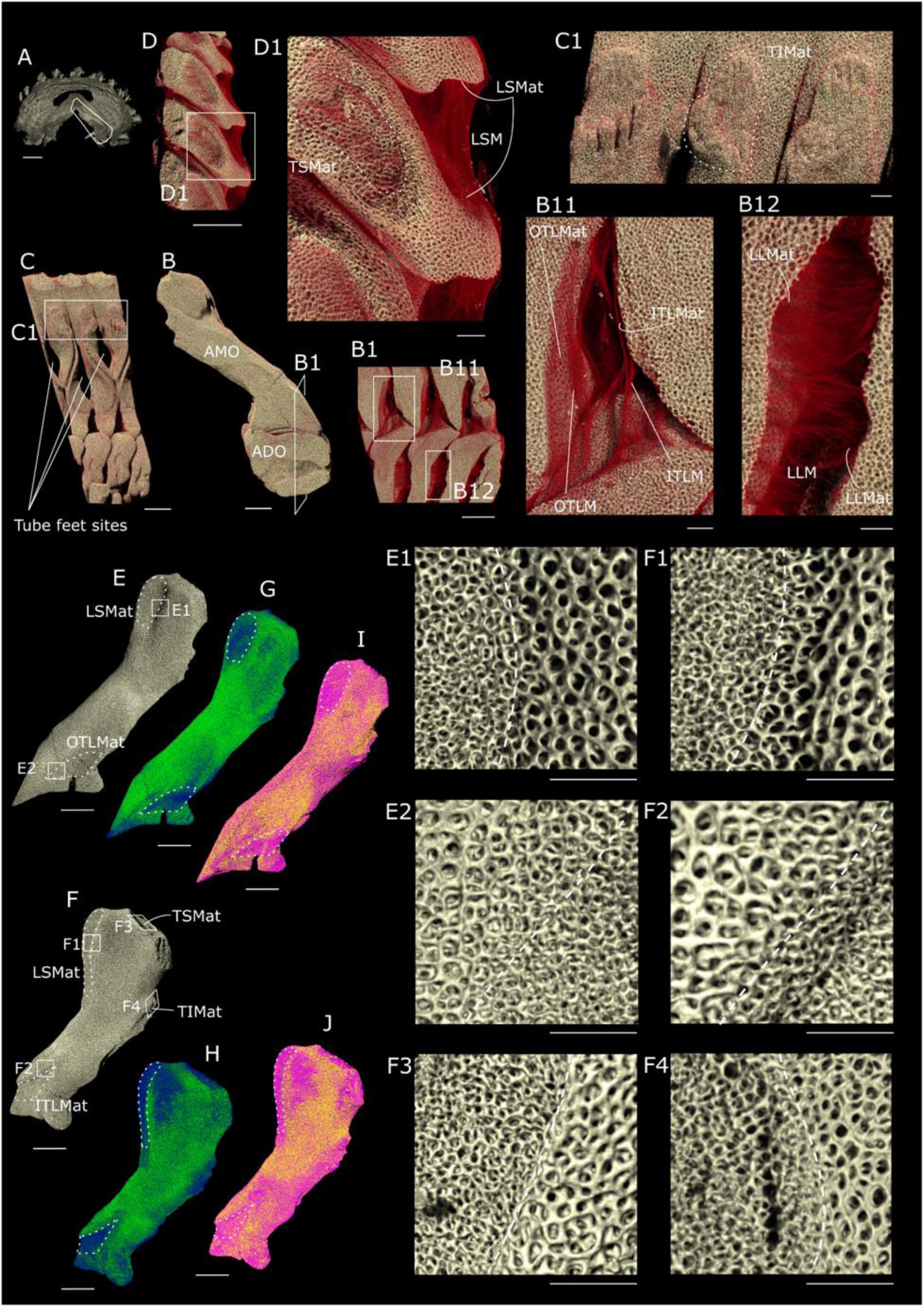
Stereom structure of ossicles at muscle and non-muscle attachment sites (**A**) Cross-sectional view of a starfish ray when observed from the proximal side. It reveals the arrangement of ambulacral (AMO) and adambulacral ossicles (ADO) forming a ridge-shaped pattern symmetrically on both sides of the ridge. High-resolution scans were conducted on ossicles from one half of the ridge. (the scan was performed at 13 µm voxel size and is adapted from schwertmann) (**B**) Geometrical arrangement of Ambulacral (AMO) and adambulacral ossicles (ADO) when visualised in a high-resolution scan from the proximal side of the ray (the scan was performed 1 µm voxel size). (**B1**) A zoomed view of the cross-section showing the connecting muscles between the ambulacral and adambulacral ossicles (AMO-ADO) and adjacent adambulacral ossicles (ADO) is shown. (**B11**) Each adambulacral ossicle (ADO) is attached with two muscle groups with ambulacral (AMO) ossicles. The outer transverse lateral muscles (OTLM) connect distal side of ambulacral ossicle (AMO) to the adambulacral ossicle (ADO), while the inner transverse lateral muscles (ITLM) connect the proximal side of the ambulacral ossicle (AMO) to the adambulacral ossicle (ADO). The sites at which, these muscle groups attach to the ambulacral ossicle (AMO) are outer transverse lateral muscles attachment site (OTLMat) and inner transverse lateral muscles attachment site (ITLMat). (**B12**) The longitudinal lateral muscles (LLM), which connect adjacent adambulacral ossicles to each other is visualised. (**C**) Left hand side view of the ambulacral (AMO) and adambulacral ossicles (ADO) assembly of image(**B**). Gaps between adjacent ambulacral ossicles (AMO) are the sites, where tube feet do exist. (**C1**) The attachment sites of transverse infra-ambulacral muscles (TIMat) are shown, and one of the attachment sites is depicted with a dotted line. Transverse infra-ambulacral muscles (TIM) link the neighbouring ambulacral ossicles situated on the opposite side of the ridge. TIM were excised to remove the ossicles from one side of the ridge and therefore not visible in the scan. (**D**) Top view of the geometrical arrangement of the ambulacral ossicles (AMO) of image(**B**). (**D1**) The attachment sites of transverse supra-ambulacral muscles (TSMat), which is a depression are visualised. Like Transverse infra-ambulacral muscles (TIM), Transverse supra-ambulacral muscles (TSM) also connect the neighbouring ambulacral ossicles situated on the opposite side of the ridge. TSM were excised to remove the ossicles from one side of the ridge and therefore not visible completely but some remains of the excised muscles could be seen at the muscle attachment site (TSMat). Longitudinal supra-ambulacral muscles (LSM), which connect adjacent ossicles next to each other on the same side of the ridge are seen. LSM connects the proximal side of one of the ambulacral ossicles (AMO) to the distal side of the adjacent one. (**E**) An individual ambulacral ossicle (AMO) is visualised from the distal side of the ray. From this view, the location of the muscle attachment sites LSMat and OTLMat (highlighted with a dotted line) can be seen on the ambulacral ossicle (AMO). (**F**) An ambulacral ossicle (AMO) is visualised from the proximal side of the ray. The view is obtained such that muscle attachment sites LSMat and ITLMat (highlighted with a dotted line) could also be visualised along with TSMat and TIMat. (**E1, E2, F1-4**) Images show the magnified view of the stereom of the inter ossicular muscle (IOM) attachment sites next to non-muscle attachment sites (a dotted line is drawn to mark the separation between two sites). It is seen that the stereom at the IOM attachment sites is finer as compared to non-muscle attachment sites. (**G**, **H**) The images show the stereom thickness distribution at the surface of an ambulacral ossicle (AMO). Stereom with thickness below 7 µm is visualized in blue, while that exceeding 7 µm is visualized in green. Notably, all inter-ossicular muscle (IOMs) attachment sites exhibit finer stereom with thicknesses below 7 µm, hence appearing in blue. While, non-muscle attachment sites showcase thicker stereom exceeding 7 µm, represented in green. (**I, J**) The images show the size distribution of porosities of the stereom at the surface of an ambulacral ossicle (AMO). Porosities with a size below 9 µm are visualized in magenta, while those exceeding 9 µm are visualized in yellow. Notably, at all muscle attachment sites stereom exhibit finer porosities with size below 9 µm, hence appearing in magenta. While, at non-muscle attachment sites, at most of the locations, stereom showcases thicker porosities exceeding 9 µm, represented in yellow. Scale bars: (**A**) 1mm; (**B, B1, C, D, E, F, G, H**) 500 µm; (**B11, B12, C1, D1, E (1&2**), **F** (**1-4**)) 100 µm

To test this hypothesis, we analyzed the thickness of the stereom at six muscle attachment sites of the ambulacral ossicle and compared it to non-muscle attachment sites of the same ossicle (Figure 3 (E, F), SV 3). An enlarged view of stereom at the muscle attachment sites next to the non-muscle attachment sites is shown in Figure 3 (E1,2, F1-4). Our data shows that at muscle attachment sites, the stereom structure is “finer” in comparison to the structure at the non-muscle attachment sites.

We also performed a 3D thickness distribution analysis of the stereom of the ambulacral ossicle. We found in comparison to the non-muscle attachment sites the average thickness of stereom at muscle attachment sites is typically less than 7 µm and the associated pore size is typically less than 9 µm (see Figure 3 (G-J), SV 4, SV 5).

### Correlation of the stereom with mechanical stresses generated by the IOMs

Mechanical load arising from muscular forces not only generates stresses in 2D at the muscle attachment sites on the surface of the ossicle, however also induce non-uniform stresses in the 3D stereom of an ossicle. If the entire stereom is correlated to these complex mechanical stresses, then the 3D distribution of von Mises stresses should also be linked to the 3D structure thickness distribution of stereom.

To test this hypothesis, we used high resolution X-ray CT data (voxel size of 1 µm) from an ambulacral ossicle and recreated the entire shape using a solid homogenous CAD geometry, without any stereom. Using finite element analysis (FEA), we then applied tensile forces at the muscle attachment sites (Figure 3 (E&F), SV 3) and determined the von Mises mechanical stress distribution within the “solid” ossicle. We then compared the FEA calculated distribution of the stresses with the stereom structure thickness distribution of the original ossicle (Figure 4).

**Figure 4:**
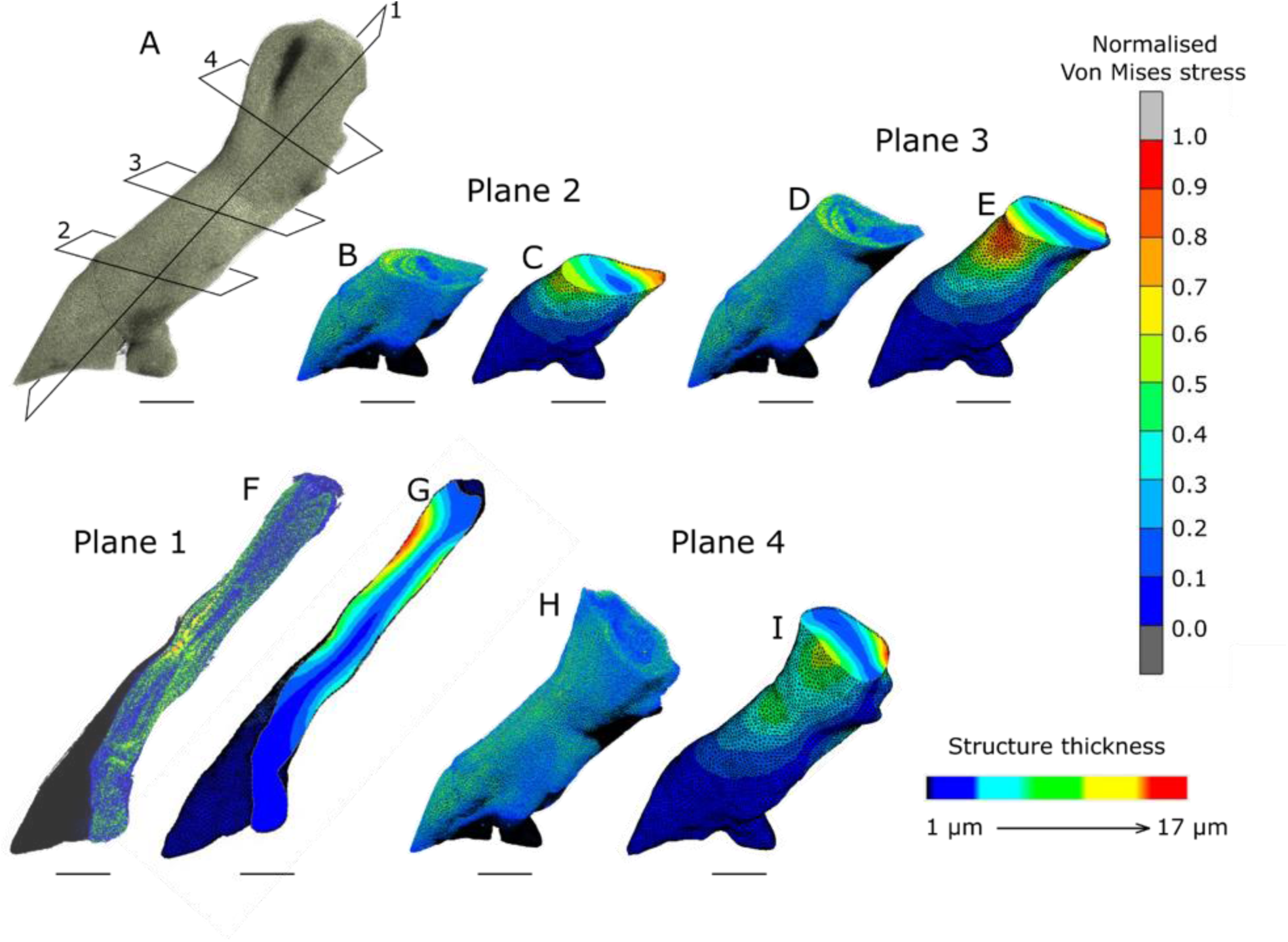
Correlation of von Mises stresses and stereom structure thickness (**A**) X-ray CT scan image of an ambulacral ossicle (AMO), with different planes drawn at various locations for comparing the thickness distributions of stereom and von Mises stress distributions at these cross-sections. (Scan was performed at 1 µm voxel size) (**B, D, F, H**) These images display the distribution of stereom structure thickness at different cross-sections of the ambulacral ossicle (AMO). The colour gradient, from blue to red, represents thickness variations, ranging from low to high. (**C, E, G, I**) These images illustrate the von Mises stress distribution at different cross-sections of the ambulacral ossicle (AMO). The colour spectrum, from blue to red, indicates the magnitude of von Mises stresses, ranging from low to high. Scale bars: (**A-I**) 500 µm

Our results show a very good agreement of the 3D von Mises mechanical stress distribution with the 3D stereom structure thickness distribution (Figure 4). The 3D stereom structure thickness distribution varies from 1 µm to 17 µm. We also observed that in regions with lower von Mises stresses the stereom typically has lower structural thickness than in regions with higher stresses. This agreement is consistent throughout the ossicle and can be observed at various cross-sections in different planes.

### Stereom at inter-ossicle contact sites

When two contact surfaces of major load-carrying ossicles touch, they exert normal forces onto each other. If the stereom correlates with the compressive forces, then the stereom at these contact sites should be different from that of the non-contact sites. To test this hypothesis, we analysed the stereom at the local contact and non-contact sites in adambulacral ossicles. Adambulacral ossicles are connected to one of the biggest IOMs and share biggest contact surfaces with each other (Figure 3 (B1, B12), Figure 5 (B)).

**Figure 5:**
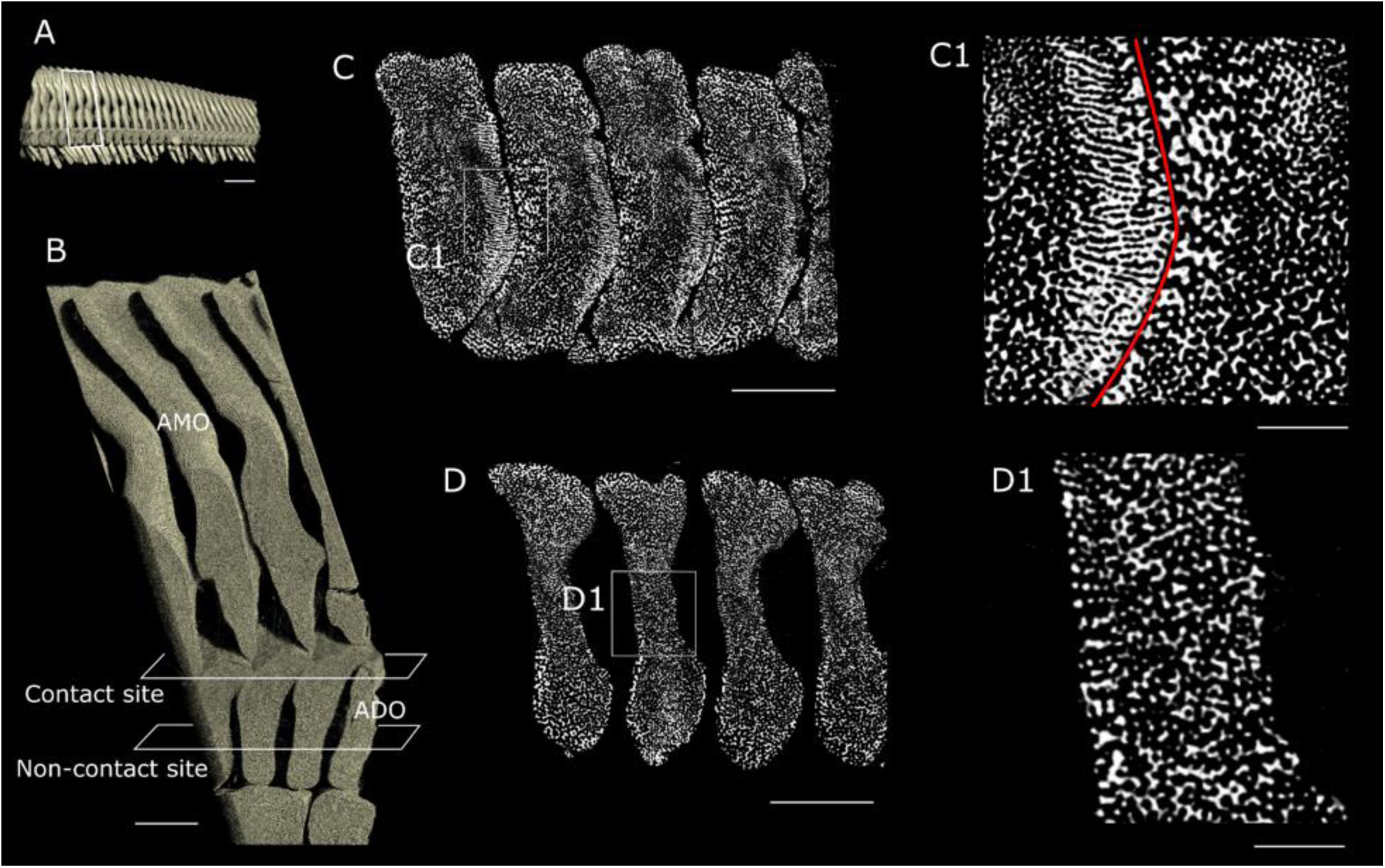
Stereom structure of ossicles under compressive stress (**A**) X-ray CT scan image of the ridge structure formed by the assembly of ambulacral (AMO) and adambulacral ossicles (ADO) (the scan conducted at a resolution of 13 µm). (**B**) This cross-sectional view provides insight into the geometric arrangement of adambulacral ossicles (ADO), highlighting the contact and non-contact regions between adjacent adambulacral ossicles (ADO). (the X-ray CT scan was performed at 1 µm resolution). (**C**) This image shows a cross-sectional view of the stereom at the contact sites, where adjacent adambulacral ossicles (ADO) share contact surfaces. In the zoomed-in view (**C1**), the red line demarcates the contact line between adjacent ossicles. At the contact site, the stereom is aligned perpendicular to the contact line. On the left side of the contact line, the adambulacral ossicle (ADO) surface is convex, with thinner and closely spaced stereom, while on the right side, the ADO surface is concave, featuring thicker and widely spaced stereom compared to the opposite side. (**D**) Here, the cross-sectional image reveals the stereom at non-contact sites. A closer look in (**D1**) shows no specific alignment of the stereom. Scale bars: (**A-D**) 500 µm; (**C1, D1**) 100 µm

Figure 5 (B) shows the sectional side view of the adambulacral ossicles assembly. In this figure, the location of contact sites between adjacent adambulacral ossicles can be seen. Figure 5 (C, D) shows the cross-section of stereom present at contact and non-contact sites respectively. The contact site between adjacent adambulacral ossicles is a curved surface and could be seen as an arc in the 2D cross-sectional view (Figure 5 (C1)). The contact surface for the adambulacral ossicle on the left side of the contact site is convex while it is concave for the ossicle on the right. Our results show that at the contact site, the stereom is oriented in the direction normal to the line of contact in both the ossicles (Figure 5 (C1)). However, the stereom on the convex surface is relatively thinner as compared to that on the concave surface. Our data shows no such alignment in stereom observed at non-contact sites of ossicles (Figure 5 (D1)).

### Growth and remodeling of ossicles

Starfish undergo two primary growth mechanisms of their ossicles. To increase the length of the rays new ossicles form at the tips throughout the lifespan of a starfish. In addition, as the starfish age, the ossicles already present in their skeleton also increase in size by adding new layers of stereom on the outside [34]. Consequently, the ray of a mature starfish is a kind of “archive” for ossicles of different ontogenetic stages. Ossicles at the base (proximal side) of the ray are the oldest and most mature structures with additional material deposited on a central nucleus region, whilst small ossicles at the tip are the youngest ossicles (Figure 6). Among all the types of ossicles, the ambulacral and adambulacral ossicles maintain consistent geometric shapes along the entire length of the ray, with a gradual variation in their sizes. To investigate whether these ossicles just grow by depositing material externally or if they also have the ability to remodel already deposited internal stereom, we conducted an analysis comparing stereom distribution patterns at the nucleus of ambulacral ossicles (AMO) across different age groups (Figure 6(A(1-3), A1(1-3)). Our assessment involved the calculation and comparison of the APP of these central nucleus regions (see Figure 6(C)). If the ambulacral ossicles grow by depositing material on the outside and are unable to remodel already deposited stereom, then a central nucleus should always be present in every grownup ossicle and the APP of this nucleus should be the same in every ambulacral ossicle.

**Figure 6:**
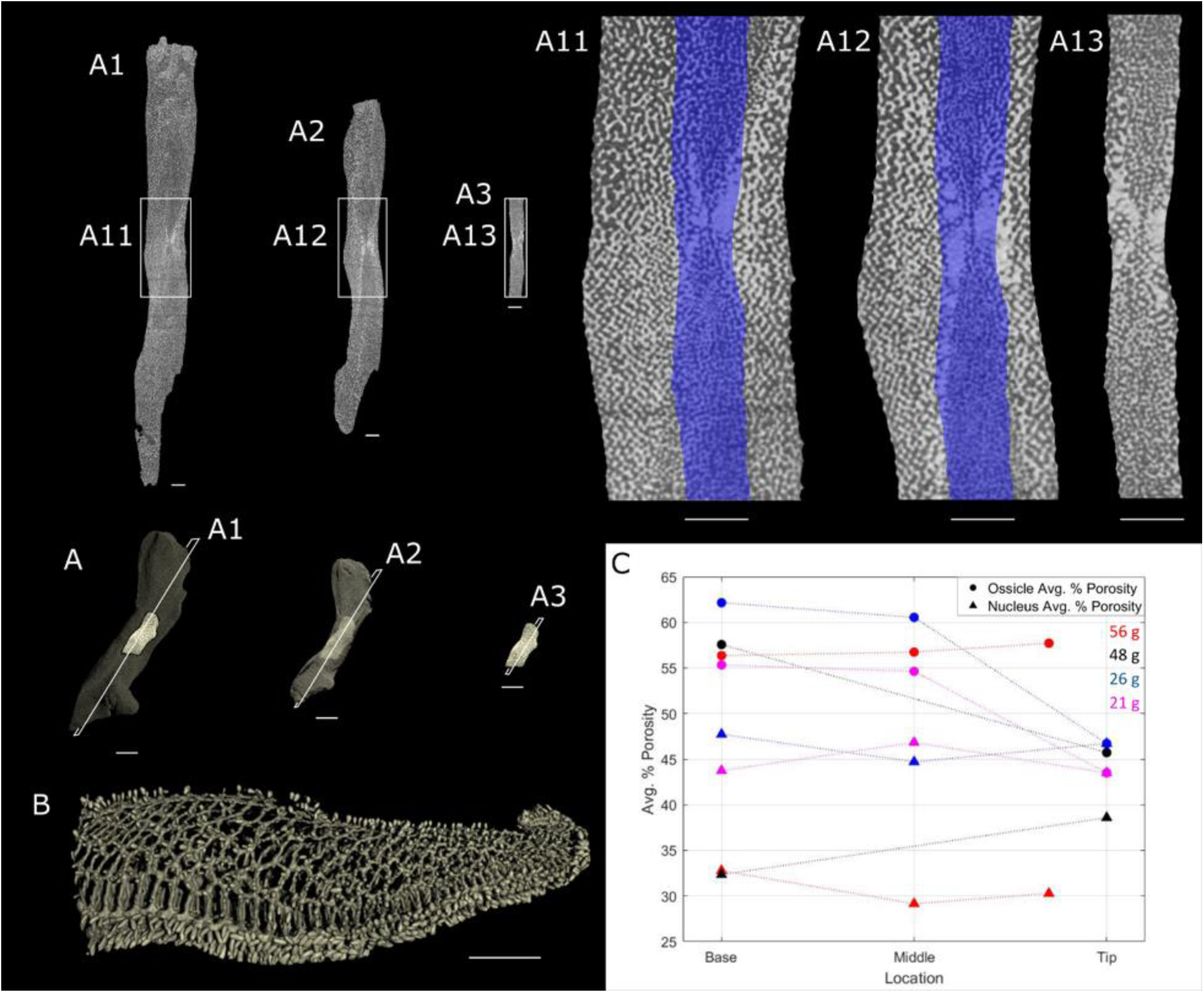
Ontogenetic changes of stereom structure (**A**) Progressive growth of ambulacral ossicles (AMO) along the length of a ray over time. The solid-colored small ossicle at the ray’s tip symbolizes the youngest ambulacral ossicle (AMO). The faded colouration of ambulacral ossicles (AMO) depicts mature ossicles in the middle and base sections of the ray. The fully coloured young ambulacral ossicle (AMO) positioned at the centre of mature ossicles supports the hypothesis that each mature ossicle is surrounded by a younger ossicle at its core. (Scans were performed at 1 µm and 2 µm voxel size) (**A**(1-3)) Stereom patterns at the cross-section of a plane passing through the centre of the ambulacral ossicles (AMO) of different ages (from base, middle and tip region). The distribution of stereom pattern is similar in all three ambulacral ossicles. **A**(11, 12, 13) Enlarged view of the stereom with geometrical shape of tip AMO ossicle (shown in blue colour) being projected over middle and base AMO. The stereom of AMOs from middle and base section show a similar distribution pattern to the stereom from the AMO from the tip section (blue shading). (**B**) Presented here is an X-ray CT scan image of a ray In conjunction with Figure (**A**), this image delineates the base, middle, and tip regions from which ambulacral ossicles (AMO) were extracted for analysis. (the scan was performed at 50 µm voxel size). (**C**) The graph illustrates a comparison between the average percentage porosity (APP) values of complete ambulacral ossicles (AMO) and their nucleus regions. Scale bar: (**A**) 500 µm; (**B**) 10 mm; (**A**(1-3)) 150 µm; (**A**1(1-3)) 150 µm

Our findings indicate that across all tested starfish, the APP in the ambulacral ossicles increases with ageing, with the lowest values observed in the youngest ossicles and the highest in the oldest ossicles. However, the APP in the central nucleus region remains relatively consistent irrespective of aging (Figure 6 (C)).

## Discussion

Both Echinoderms and Verterbrates possess a mesodermal endoskeletons. Both vertebrate bones and echinoderm ossicles share a porous microstructure. In vertebrates, bones have demonstrated a clear correlation with mechanical loading and the ability to remodel their porous microstructure in response to changes in mechanical stresses. This adaptation ensures that the bone structure remains optimized for its mechanical demands.

One of the key differences between these two types of endoskeletons is however their respective growth pattern. Echinoderms increase the size of their skeletons by adding new ossicles over time and by enlarging previously formed ossicles. It’s not clear whether echinoderms possess a similar capacity for remodeling these ossicles in response to mechanical stress. If echinoderms do indeed exhibit this capability, it would seem likely that the response to mechanical stress and the capacity for remodelling are fundamental traits of mesodermal endoskeletons which of both echinoderms and vertebrates. If not, then this trait would not apply universally to all mesoderm endoskeletons, emphasizing the diversity of adaptive mechanisms in different phylogenetic branches.

In our research, we conducted a comparative analysis of the stereom in various ossicle types across different age groups. Our findings indicated that the APP of the stereom differs significantly depending on both the ossicle type and its age (Table 1). Specifically, ambulacral ossicles were found to have the lowest APP (Figure 2) and thus exhibit the most solid structure among all the different ossicle types we examined. It’s worth noting that ambulacral ossicles are not only the most solid but also the largest in size compared to other ossicle types. Additionally, they are connected to the most extensive network of inter ossicular muscles (IOMs).

Subsequently, we selected the ambulacral ossicles for a comprehensive and detailed structural analysis. In our investigation, we compared the 3D stereom thickness distribution to the 3D von Mises mechanical stress distribution and identified a significant correlation between the two (Figure 4). Wherever mechanical stresses are higher within the ambulacral ossicle, we observed the presence of thicker stereom. This thicker stereom serves as a structural adaptation to effectively counteract and manage the increased mechanical stresses, ensuring the ossicle’s ability to withstand these stresses. Conversely, in regions where mechanical stresses were lower, we observed thinner stereom. This thinner stereom structure in low-stress regions points towards a more efficient allocation of resources within the ossicle, potentially increasing the efficiency of material use. Our results strongly suggest that the variation in stereom thickness across the ossicle is not a random occurrence; rather, it directly correlates with the mechanical stresses present in those specific regions. This correlation underscores the adaptive nature of the ossicle’s stereom structure in response to mechanical demands.

In our analysis, we specifically examined the stereom at the local sites experiencing high mechanical stress, such as muscle attachment sites. Our findings revealed that at the muscle attachment sites of an ambulacral ossicle, the stereom appears to be finer and has smaller porosities as compared to the stereom at non-muscle attachment sites (Figure 3, SV 3, SV 4, SV 5). For a given surface area such a structure provides a greater number of attachment sites for muscle fibers as compared to thicker stereom with bigger porosities, which would result in a better distribution of mechanical stresses over the surface. As a consequence, a reduction in stress concentration at these sites would contribute to enhanced mechanical stability and resilience. This adaptation reflects the efficiency of the ossicle’s structure in responding to specific mechanical demands.

The other local sites where stereom is subjected to high mechanical stresses are the locations where two ossicles share contact surfaces and transmit mechanical loads. One of the biggest contact sites is shared among adjacent adambulacral ossicles. At these local contact sites, we found that the stereom orients itself in the direction normal to the contact surface (Figure 5). The type of load being transmitted through these contact surfaces is the normal compressive load. The geometrical alignment of the stereom in the direction of mechanical load allows the local stereom to undergo compressive normal stresses. This is particularly valuable since ossicles are composed of brittle magnesium calcite, a material well-suited to endure compressive stresses in contrast to tensile and shear stresses [24]. This alignment and geometry of the stereom at these local contact sites thus demonstrate a clear correlation with the mechanical load and ultrastructure.

Ossicles at various locations along the length of a starfish ray serve distinct functions and thus face different types of mechanical loads. For instance, during feeding, the tube feet located at the base and middle segments of the ray are responsible for prying open mussels, placing direct loads on the attached ambulacral ossicles. In contrast, the ambulacral ossicles at the tip of the ray are not subject to such loads [3]. During ontogenesis, new ossicles are continually added at the ray’s tip, lengthening it. Consequently, the older ossicles that were once at the tip now form the middle and base regions, resulting in a shift in their functionality. Different stereom structures may be better suited for such changing mechanical demands. If starfish possessed the ability to dynamically alter their previously deposited stereom, we would expect a probably entirely different stereom structure throughout the ossicle. However, our findings indicate otherwise. The distribution patterns of stereom at the nucleus of mature ambulacral ossicles (AMO) closely resemble those observed in newly formed ossicles, indicating the lack of stereom structure remodeling with aging (see Figure 6(A11, A12, A13)). Interestingly, our observations reveal that, for every starfish the APP in the central nucleus region of ambulacral ossicles (AMOs) consistently was lower compared to the APP of the entire ossicles. This discrepancy is attributed to the presence of a thicker stereom in the nucleus region of these ossicles (Figure 6(A11, A12, A13)). Remarkably, the APP value of the nucleus region was relatively constant across ossicles of different age groups, suggesting no stereom remodeling occurs in the central nucleus region over time, despite changes in mechanical stress distribution within the ossicle (see Figure 6(C)).

During our analysis, we also found a single retinal ossicle which presumably was fractured during some time in the life of the starfish. Interestingly, the “healed” osscile apparently did not undergo the remodelling of the broken overlapping surfaces. Instead, a patch-up or repair was evident. If ossicles possessed the capability to remodel the stereom, we would expect to see a seamless continuity in the stereom at the cracked site, rather than a patch-up (SF 2). This result, along with our previous one, supports our findings that starfish may not possess the ability to dynamically remodel the stereom that has already been deposited.

## Conclusion

Our study clearly shows that the ultrastructure of starfish ossicles is closely correlated to the mechanical stresses they encounter. While this ability to adapt to mechanical stresses is well-documented in vertebrate bones, our research suggests that it may be a general feature of the mesoderm endoskeleton in the Deuterostomia taxonomic group. However, our results also show that not all ossicles in the echinoderm endoskeleton have the ability to remodel their structure depending on the change of mechanical stress. Consequently, the remodeling of endoskeleton material could be a specific trait or ability unique to mesodermal vertebrate endoskeletons.

## Materials and Methods

### Specimen and sample preparation

Specimens of the starfish *Asterias rubens*, were provided by the Alfred Wegener Institute in Helgoland. We selected four different individuals representing different ontogenetic stages with body mass ranging from (21 – 56) g.

Following established protocols, the starfish specimens were relaxed in freshwater and subsequently fixed in a solution consisting of 4% formaldehyde and seawater sourced from the North Sea. After fixation, the specimens were permanently preserved in a solution of 4% formaldehyde and distilled water. Following preservation, they were transported to the University of Applied Sciences in Bremen, Germany. On site, the five different types of ossicles ambulacral, adambulacral, marginal, reticular and carinal were carefully extracted from the base, middle and tip regions of the starfish ray (Figure 2 (A)). Ossicles where then placed in small plastic containers, stabilized with tissue and scanned in air.

### X-Ray Microtomography

The overview scan of complete starfish skeleton was performed at 76 µm voxel size with a Phoenix micro-CT scanner (v|tome|x m research edition, GE Digital Solutions, 130 kV and 100 µA, 2024 projections, 0.33 s exposure time). The associated reconstruction software (Phoenix dato|x 2, GE Digital Solutions, Boston, USA) was used for the post-processing of the scans. The scan was adapted from [35].

The overview scan of the complete starfish ray was performed at 50 µm voxel size with Bruker Skyscan 1275 X-ray micro CT scanner using X-rays, 60 kV and 61 µA, 1056 projections, 0.1 s exposure time). Bruker NRecon software was used for reconstruction.

Selected regions of the of starfish ray (Figure 1:Figure 1_(B-E)) were scanned at 13 µm voxel size with ZEISS Xradia Versa 520 XRM machine, 70 kV and 86 µA, 3201 projections, 1.5 s exposure time and 4x optical magnification). ZEISS Reconstructer Scout-and-scan software was used for reconstruction. This specific scan data was re-used from [35].

The high-resolution X-Ray CT scans of individual ossicles were performed on ZEISS Xradia Versa 520 XRM machine, 80 kV and 88 µA, 3000 projections, 4x optical magnification). For these scans five seconds of exposure time per projection were used for high contrast. ZEISS Reconstructer Scout-and-scan software was used for reconstruction.

’Initially, we performed a scan of an ossicle at a voxel size of 1 µm and calculated the APP for the ossicle. Subsequently, we digitally processed the CT image dataset to reduce it to a 2 µm voxel size and recalculated the APP. The APP difference between the 1um and 2um scans was only around 1 %. To notably reduce scan time we thus performed APP related scans of single ossicles using a 2 µm voxel size.

However, the calculations for 3D structure thickness distribution are highly sensitive to changes in pixel size. Consequently, we conducted scans at a 1 µm voxel size for these specific calculations. Table 2 summarize all the scan parameters for above mentioned X-Ray CT scans.

**Table 2:**
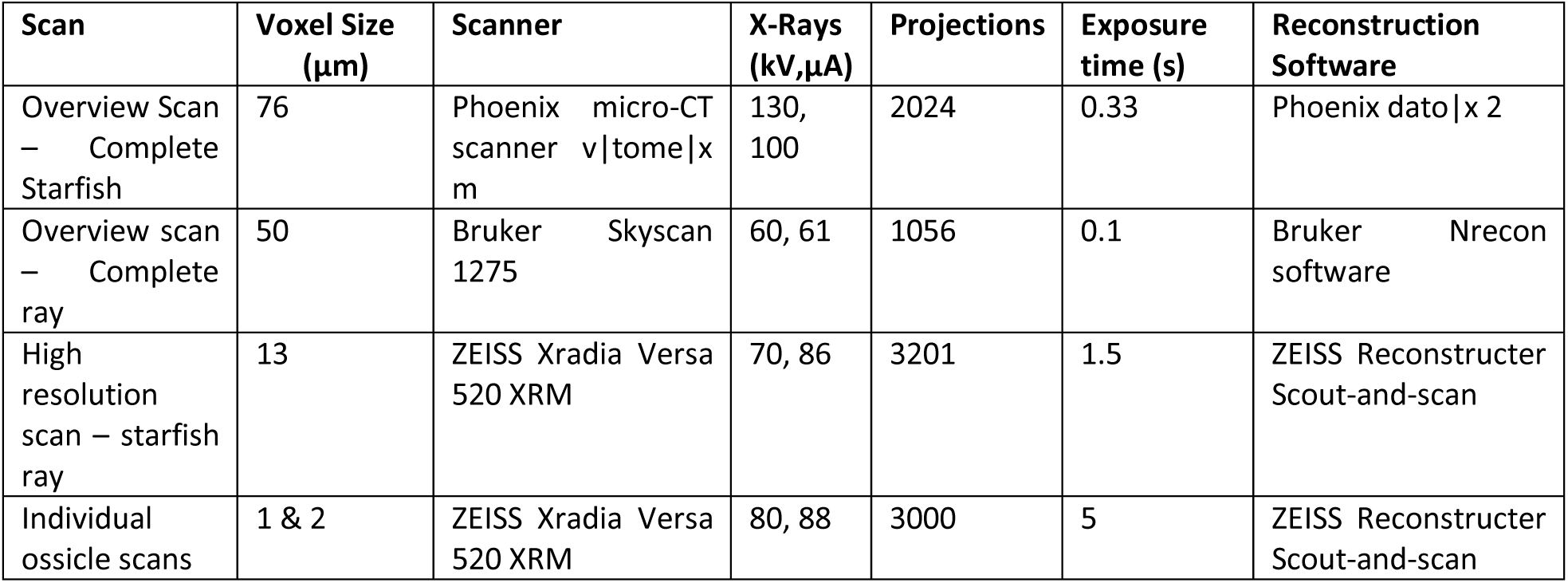
A summary of scan parameters for different X-Ray CT scans that have been used in this study.

### Processing of 3D X-Ray Image Data

Image processing and visualization was carried out using Bruker CTAn software, while the visualization of the 3D CT data was accomplished using volume rendering software Bruker CTVox.

Adaptive thresholding (mean of the maximum and minimum values), was employed to establish a threshold for the ossicle stereom, effectively distinguishing it from the background. This method is used to dynamically adjust the threshold for image segmentation based on the local characteristics of the image, ensuring an accurate separation of the trabecular structures from the surrounding background.

To calculate the percentage of volume inside a porous ossicle, we generated a solid-filled representation of the ossicles and subsequently subtracted the actual stereom geometry from it, as illustrated in (SF 1). The average percent porosity (APP) was determined using the following formula:

APP = (Total volume of solid-filled ossicle - Total volume of stereom)/ (Total volume of solid-filled ossicle) * 100%

This calculation method allowed us to quantify the porous volume within the ossicles.

True 3D structure thickness calculations for CT image data were performed using a model-independent method developed by [36]. This method provides an approach to measure the thickness of structures within the 3D context of CT image data without relying on specific models or assumptions, making it a valuable tool for accurate and unbiased structural analysis [36,37].

For starfish of body mass 21g and 26g, the central nucleus region was chosen to be the complete volume of the ambulacral ossicles (AMO) form the tip. While for the starfish of body mass 48g and 56g, a central nucleus region was manually defined.

### Finite Element Analysis

A CAD geometry of homogeneous solid-filled ambulacral ossicle was derived from the 3D X-ray CT scan data and used to make a finite element model (FE model). A uniform mesh with linear isoparametric three-dimensional tetrahedron elements was used to form the ossicle geometry consisting of 123,203 elements. Material property of magnesium calcite ((Young’s modulus) E = 100 GPa) was used as calculated experimentally by [23]. The value of Poisson’s ratio was chosen to be 0.3. Pulling forces were applied on all the muscle attachment sites in the direction of respective muscle fibers. The magnitude of the resulting forces was decided by the size of the muscle attachment site. As, the number of mesh nodes present on each muscle attachment site are proportional to the surface area of the muscle attachment site. Therefore, the forces on each muscle attachment site were kept proportional to the number of mesh nodes leading to a factor in the calculated normalised von Mises stress.

### Statistics

All statistical tests were performed using R Studio software (RStudio, 2020). Data was tested for homogenity of variance using Shapiro-Wilk tests and normal distribution using Barthlett tests. If these criteria were met, ANOVA tests were performed to test for significant effects with TukeyHSD post-hoc tests to test for significant differences between groups.

## Supplementary Figures

**SF 1.**
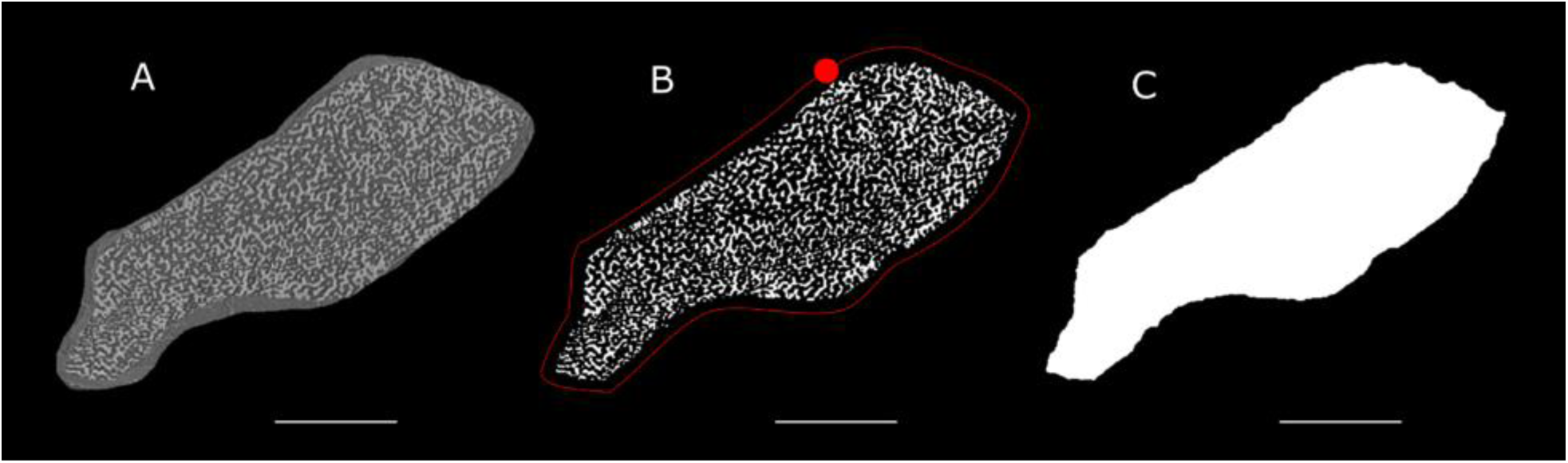
Schematic illustration of ossicle geometry analysis (A) Reconstructed slice derived from X-ray CT data of an ambulacral ossicle, showcasing the stereom in grayscale. The scan was performed at 1 µm voxel size. (B) Employing adaptive thresholding, the stereom is effectively separated from the background, transforming the image into a binary format. A ball is then utilized to roll across the binary data’s surface, creating an envelope that encapsulates the ossicle. (C) The interior volume encompassed by the envelope generated in image (B) is filled, resulting in a solid-filled representation of the ossicle’s geometry. Scale bar: 250 µm

**SF 2.**
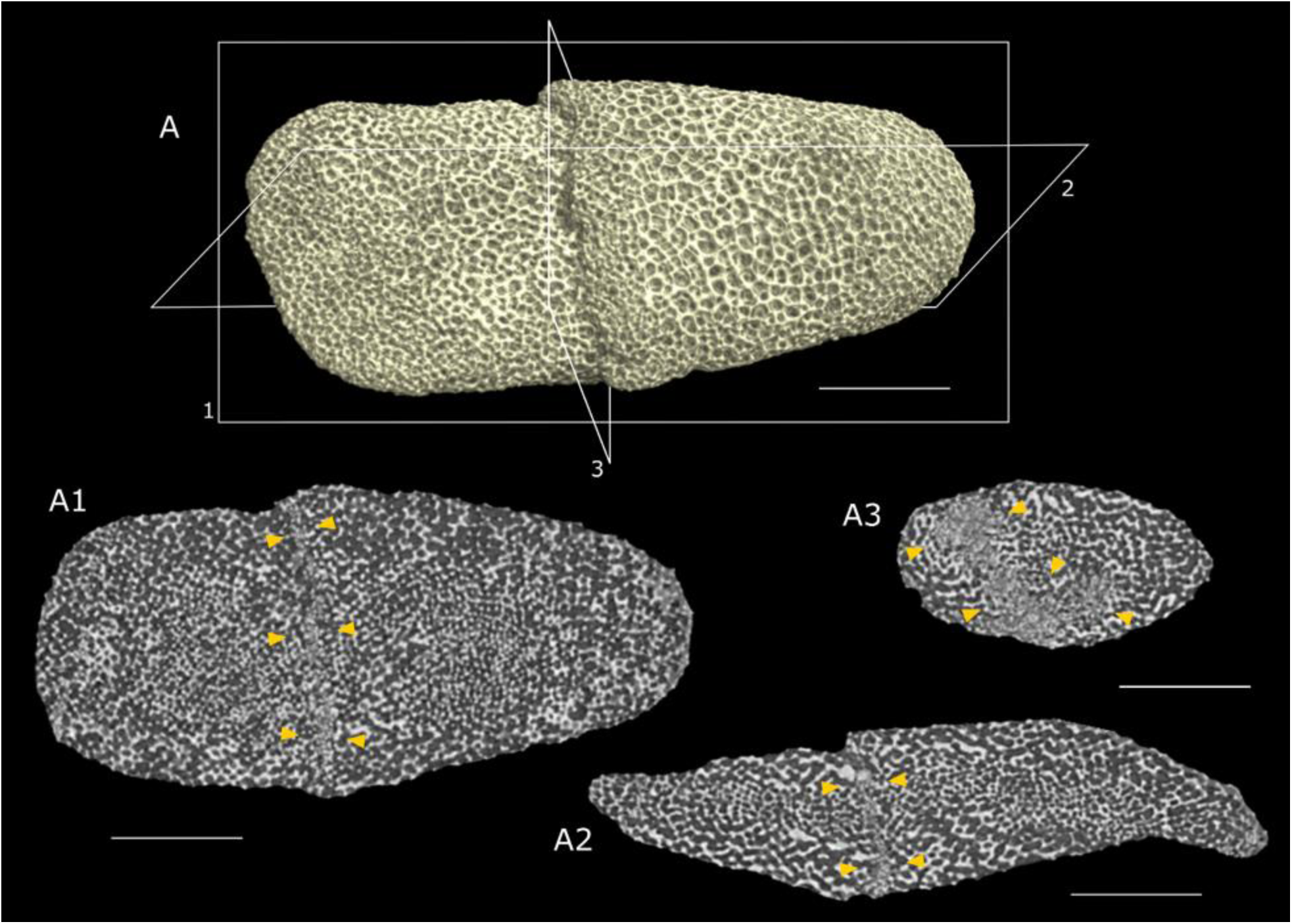
Retinal ossicle with a healed fractured surface. (A1, A2, A3) Stereom at three mutually perpendicular cross-sections of the ossicles is shown. All these cross-sections Pass through the fracture surface. In these cross-sections, the discontinuity in the stereom at the fractured site can be observed. (The X-ray CT scan was performed at 2 µm voxel size). Scale bar: 250 µm.

## Supplemental Movies

### SV 1 (separate file)

*Exploration of the intricate ossicular network within a starfish endoskeleton through captivating visualizations. Crafted using Bruker CTVox software, these visuals stem from an X-ray CT scan at 76 µm voxel size, as adapted from* [35]*. The carefully chosen colour scheme employs blue for lower greyscale values, indicating less dense structures, and green for higher greyscale values, highlighting more densely packed structures. Dive into the fascinating world of starfish anatomy with this mesmerizing representation*.

### SV 2 (separate file)

*Visualization of stereom distribution within an ambulacral ossicle using Bruker CTVox software. The visuals are derived from a high-resolution X-ray CT scan with a 1 µm voxel size capturing the complete distribution of stereom throughout the ossicle geometry. Variations in stereom geometry at different cross-sections could be observed throughout the length of the ossicle*.

### SV 3 (separate file)

*Visualization of the assembly of ambulacral (AMO) and adambulacral ossicles (ADO) accompanied by inter-ossicular muscles (IOMs) using Bruker CTVox software. The visuals are derived from a high-resolution X-ray CT scan performed at a 1 µm voxel size. The color scheme was generated by assigning the red color to the greyscale values belonging to the soft tissues and cyan color to the hard tissue (stereom). Visualisation through various cross-sections reveals the geometrical alignment and attachment sites of different IOMs. Additionally, some remains of tube feet could also be observed attached to the ambulacral ossicles*.

### SV 4 (separate file)

*Stereom thickness distribution at the surface of an ambulacral ossicle (AMO). Stereom with thickness below 7 µm is visualized in blue, while that exceeding 7 µm is visualized in green. Notably, all muscle attachment sites exhibit finer stereom with thicknesses below 7 µm, hence appearing in blue. Conversely, non-muscle attachment sites showcase thicker stereom exceeding 7 µm, represented in green. The image data from an X-ray micro-CT scan performed at 1 µm voxel size was used for this analysis*.

### SV 5 (separate file)

*Size distribution of porosities of the stereom at the surface of an ambulacral ossicle (AMO). Porosities with a size below 9 µm are visualized in magenta, while those exceeding 9 µm are visualized in yellow. Notably, at all muscle attachment sites stereom exhibit finer porosities with size below 9 µm, hence appearing in magenta. While at non-muscle attachment sites, in most of the places, stereom showcases thicker porosities exceeding 9 µm, represented in yellow. The image data from an X-ray micro-CT scan performed at 1 µm voxel size was used for this analysis*.

