## Supplementary figures and images for "The ultrastructure of the starfish skeleton is correlated with mechanical stress"

### SF 1

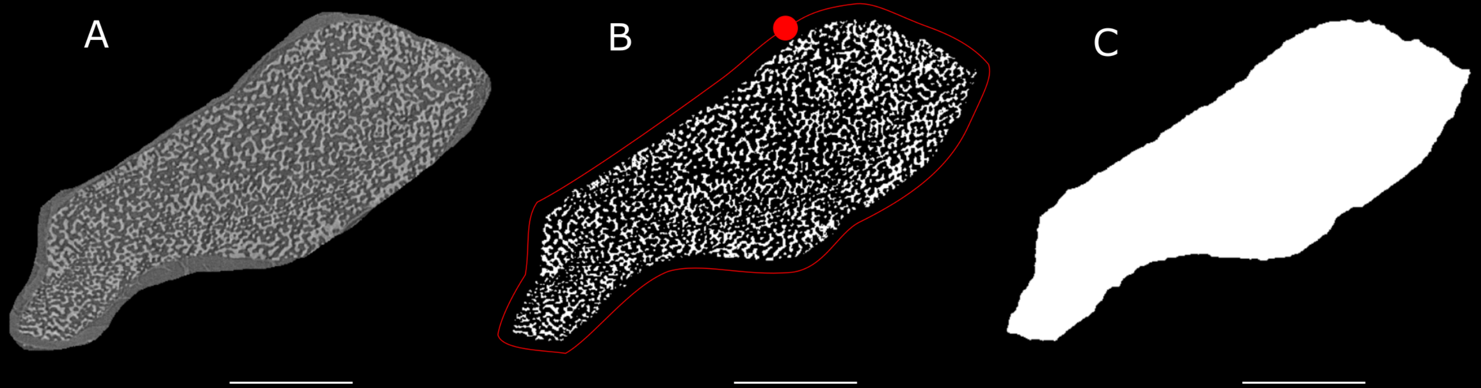

### SF 2

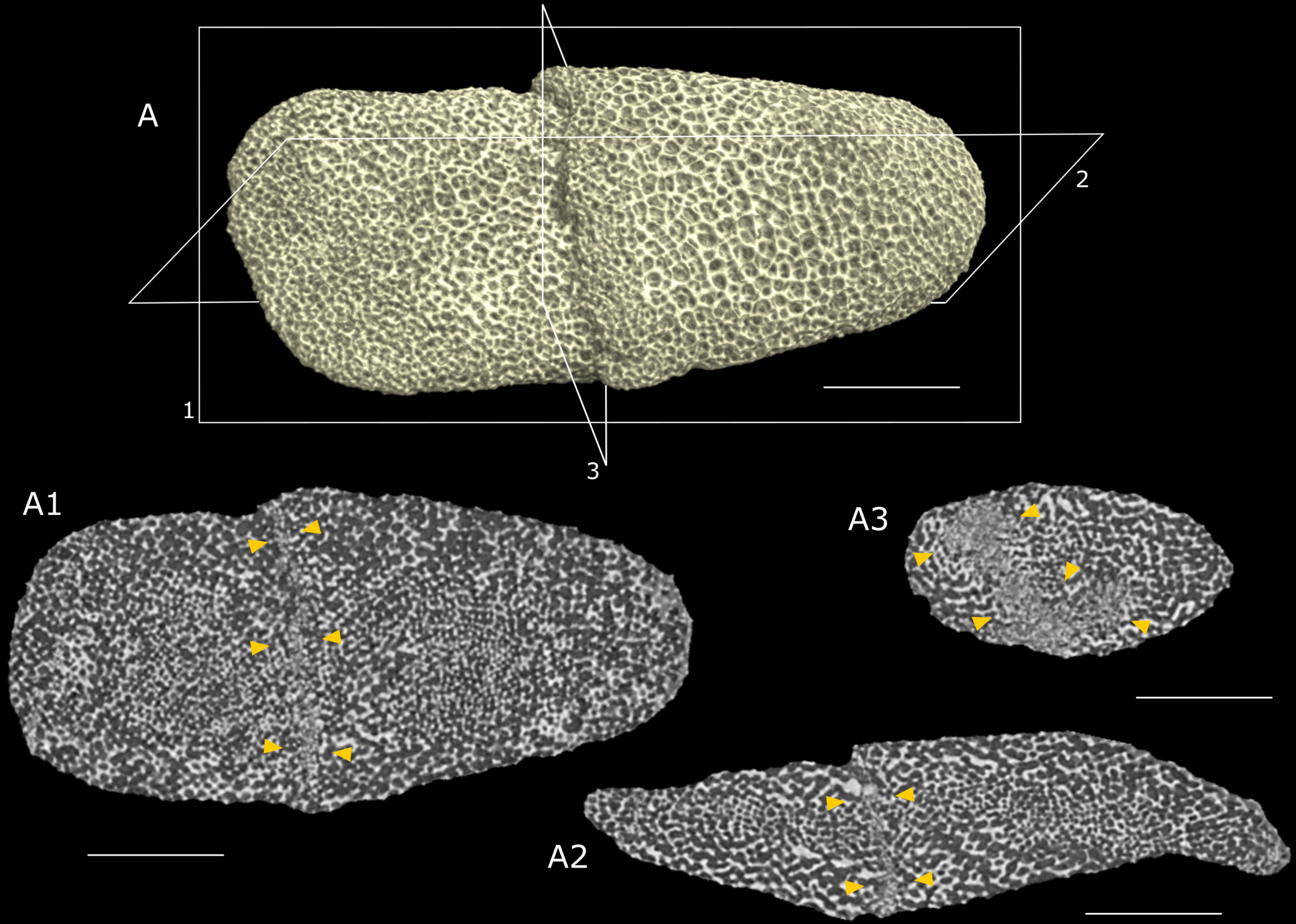
